# The astrocyte-enriched gene *Tmem44* regulates circadian protein translation in mouse astrocytes

**DOI:** 10.64898/2026.05.07.723448

**Authors:** Ji Eun Ryu, Minsung Park, Hyun Woong Roh, Jae-Hyung Lee, Eun Young Kim

## Abstract

Astrocytes contain cell-autonomous circadian clocks, but how astrocyte-enriched clock-controlled genes feed back onto the circadian clock remains poorly understood. Here, we identify *transmembrane protein 44* (*Tmem44*) as an astrocyte-enriched circadian transcript whose protein product regulates clock protein abundance through translational control. *Tmem44* mRNA oscillated in cultured mouse cortical astrocytes in a BMAL1-dependent manner, whereas TMEM44 protein was constitutively expressed and localized to the endoplasmic reticulum. Acute *Tmem44* knockdown reduced BMAL1 and PER2 protein levels without altering their mRNA levels or degradation kinetics, and dampened the amplitude and advanced the phase of *Per2*-luciferase circadian rhythms. SUnSET assays revealed that *Tmem44* knockdown decreased global nascent protein synthesis. Proximity labeling and co-immunoprecipitation further showed that TMEM44 associates with ribosomal proteins and ER-associated translational machinery. Importantly, global protein synthesis exhibited circadian oscillation in synchronized astrocytes, and this rhythmic translation was abolished by *Tmem44* knockdown. These findings identify TMEM44 as an astrocyte-enriched, ER-resident translational regulator that supports rhythmic protein synthesis and sustains proper amplitude and phase of the astrocyte circadian clock.

## Introduction

The circadian clock is a cell-autonomous timekeeping mechanism that drives approximately 24-hour rhythms in gene expression, physiology and behavior. At the molecular level, circadian rhythms are generated by core transcription–translation feedback loops (TTFLs) in which the transcription factor complex CLOCK-BMAL1 drives expression of its own repressors, *Period* (Per1/2) and *Cryptochrome* (Cry1/2), which in turn inhibit CLOCK-BMAL1 activity. A secondary stabilizing loop, in which CLOCK-BMAL1 induces *Nr1d1* and *Rorα*, whose protein products repress and activate *Bmal1* transcription, respectively, provides additional stability and precision to the oscillation ^1,2^.

Astrocytes, the most abundant glial cells in the mammalian brain, harbor cell-autonomous circadian clocks and constitute an active component of brain circadian network ^3,4^. Astrocyte circadian clocks regulate diverse aspects of brain physiology, including neurotransmitter clearance, metabolic support of neurons, and synchronization of local neural circuits ^5,6^. To systematically investigate how the circadian clock regulates astrocyte physiology, we recently conducted a circadian transcriptome analysis of cultured mouse cortical astrocytes, identifying 412 oscillating transcripts and demonstrating that rhythmic *Herpud1* expression underlies circadian regulation of ER Ca^2+^ responses ^7^.

The robustness of circadian oscillations depends not only on core TTFL components but also on accessory factors that modulate feedback loop dynamics. Rhythmically expressed non-core clock genes have been shown to modulate TTFL dynamics by influencing circadian period, phase, and amplitude in various cell types ^8–11^. However, whether cell-type specific circadian transcripts contribute to TTFL regulation in astrocytes remains poorly understood. Given that cell-type enriched transcripts are often associated with specialized cellular functions ^12^, rhythmically expressed astrocyte-enriched genes may represent previously unrecognized modulators of astrocyte circadian function.

An emerging layer of circadian regulation operates at the level of protein synthesis. Global translation has been shown to oscillate in a circadian manner in mouse liver and cultured fibroblasts ^13–15^, with a substantial fraction of rhythmically translated transcripts lacking corresponding oscillations at the mRNA level ^16–18^. This rhythmic translational control is thought to contribute to the temporal gating of protein expression and to amplify the oscillatory output of circadian clock ^13,16,19^. Nevertheless, the mechanisms that link the circadian clock to rhythmic translational activity, particularly in the brain and in non-neuronal cell types such as astrocytes, remain largely unexplored.

Here, we report that *Tmem44* is an astrocyte-enriched circadian transcript whose encoded protein is required for circadian regulation of protein synthesis. Knockdown of *Tmem44* reduced the BMAL1 and PER2 protein abundance without altering their mRNA levels or protein stability. Mechanistically, TMEM44 supports global protein synthesis and associates with ribosomal proteins at the ER. Furthermore, *Tmem44* knockdown abolished circadian oscillation of translation, accompanied by reduced clock protein abundance and dampened Per2-luciferase rhythms. Together, these findings identify TMEM44 as an ER-resident regulator of circadian translation in astrocytes and suggest that ER-associated translational control sustains the amplitude of the astrocyte circadian clock.

## Materials and Methods

### Animals

All animal procedures were approved by the Institutional Animal Care and Use Committee (IACUC) of Ajou University School of Medicine. All mice were housed in a specific pathogen-free facility at the Animal Research Center of Ajou University Medical Center under a standard 12-hour light/dark cycle, with ad libitum access to food and water.

*Bmal1*^+/-^ mice (B6.129-*Bmal1*tm1Bra/J, stock #009100; Jackson Laboratory (USA)) were initially obtained from Jackson Laboratory ^20^. *Per2*-luciferase reporter mice (B6.129S6-Per2tm1Jt/J, stock #006852; Jackson Laboratory (USA)) were obtained from Jackson Laboratory ^21^. *Tmem44*^+/-^ mice (C57BL/6N-*Tmem44*em1 (IMPC) KMPC) were produced and provided by the Korea Mouse Phenotyping Center (KMPC, Republic of Korea).

For primary astrocyte culture, postnatal day 1 (P1) littermate pups were used. For *Bmal1* experiments, *Bmal1*^-/-^ mice were used, and wild-type littermates (*Bmal1*^+/+^) were used as controls. For *Tmem44*, both wild-type (*Tmem44*^+/+^) and heterozygous (*Tmem44*^+/-^) littermates were used as controls.

In experiments using only wild-type astrocytes, P1 C57BL/6 mice were purchased from Koatech Inc. (Republic of Korea).

For brain tissue collection, adult male C57BL/6 mice (7–8 weeks old) were entrained to a 12-hour light/dark cycle for 2 weeks and then transferred to constant darkness. Mice were sacrificed on the third day of constant darkness at the indicated circadian time (CT), and the cerebral cortex was dissected.

### Identification of astrocyte-enriched circadian transcripts

Publicly available transcriptome datasets were used to identify astrocyte-enriched circadian transcripts. To assess expression in adult astrocytes, RNA-seq data from mouse cortical astrocytes were obtained from ^22^ and transcripts with mean FPKM > 10 in visual and motor cortex astrocytes were retained. To evaluate cell type–specific enrichment, a cell type–resolved cortical transcriptome dataset was obtained from ^23^.

For the Zhang et al. dataset, an astrocyte enrichment score was calculated by dividing the expression level in astrocytes by the mean expression across six major neural cell types. Transcripts within the top 10% of this distribution were defined as astrocyte enriched.

These criteria were applied to the previously identified oscillating transcripts ^7^ to obtain a refined set of astrocyte-enriched circadian candidates. Circadian rhythmicity was analyzed using the MetaCycle package in R, applying the Meta2d algorithm with default parameters. Meta2d p values were used to assess statistical significance of rhythmicity ^24^.

### Primary astrocyte culture

Primary cortical astrocytes were prepared from postnatal day 1 (P1) mouse pups as previously described ^7^. Briefly, cerebral cortices were dissected, dissociated in MEM-based medium supplemented with 10% FBS, penicillin-streptomycin, HEPES, and GlutaMAX, and plated in T75 flasks (1 pup/flask). Cells were cultured at 37 °C in a humidified 5% CO₂ incubator for 2 weeks, followed by a 7-day serum-free incubation. Astrocytes were then detached with 0.25% trypsin and plated at a density of 1.0 × 10⁶ cells per 60-mm dish for subsequent experiments.

### siRNA transfection, circadian synchronization, and bioluminescence recording

Astrocytes were seeded at a density of 1.0 × 10⁶ cells per 60-mm dish. When cells reached >90% confluency, the medium was replaced with serum-free MEM. After 72 hr under serum-free conditions, siRNA was transfected using Lipofectamine RNAiMAX (Thermo Fisher (USA), 13778150) according to the manufacturer’s instructions. ON-TARGETplus SMARTpool siRNAs were purchased from Dharmacon (USA): non-targeting control siRNA (D-001810-01-50) and *Tmem44* siRNA (L-060116-01-0005).

For circadian bioluminescence recording, primary astrocytes were prepared from *Per2*-luciferase reporter mice. When cells reached >90% confluency, the medium was replaced with serum-free MEM. After 72 hr under serum-free conditions, circadian synchronization was induced by replacing the medium with MEM containing 50% horse serum for 2 hr ^25^. Following synchronization, cells were washed twice and maintained in recording medium consisting of phenol red-free MEM (Sigma-Aldrich, M3024) supplemented with 10 mM HEPES, 4 mM NaHCO₃, 1X GlutaMAX, 1X penicillin–streptomycin, and 400 µM D-luciferin, and bioluminescence was continuously recorded using a LumiCycle luminometer (Actimetrics, USA).

Circadian time was defined based on time after serum shock (TASS), with TASS 8 hr corresponding to CT0 ^7^. LumiCycle data were baseline-subtracted using LumiCycle Analysis software, and circadian parameters including period, phase, and amplitude were calculated using the BioDare2 web tool ^26,27^ with MESA (maximum entropy spectral analysis) fitting.

### Quantitative reverse transcription polymerase chain reaction (qRT-PCR)

Total RNA was extracted from cells and purified using the RNeasy Plus Micro Kit (Qiagen, Germany, 74034). 1 μg of total RNA was reverse transcribed using an oligo-dT primer and PrimeScript RTase (TaKaRa (Japan), 2680A). Quantitative real-time PCR was performed using a Rotor Gene Q (Qiagen) with TB Green Premix Ex Taq (Takara, RR420A). The specific primers used were provided in Supplementary Table 3. SDHA was used as the reference gene for normalization in all experiments. Data were analyzed using Rotor-Gene 6000 software and quantified by the 2<–ΔΔCt method, where ΔΔCt = [(Ct__target_ – Ct__reference_) of experimental group] – [(Ct__target_ – Ct__reference_) of control group].

### Immunoblotting

Astrocytes were lysed using modified RIPA buffer (50 mM Tris-HCl pH 7.4, 1% NP-40, 0.5% sodium deoxycholate, 150 mM NaCl) supplemented with protease inhibitor cocktail (Sigma-Aldrich (USA), P8340) and phosphatase inhibitor cocktails 2 and 3 (Sigma-Aldrich, P5726 and P0044). Proteins were separated by SDS-PAGE and transferred to polyvinylidene fluoride (PVDF) membranes. After blocking with 5% skim milk, membranes were incubated overnight at 4 °C with primary antibodies at the following dilutions: anti-*Bmal1* (Abcam (UK), ab93806), 1:2000; anti-*Tmem44* (this study, #12), 1:1000; anti-Per2 (Alpha Diagnostic Intl. (USA), PER21-A), 1:1000; anti-Vinculin (Sigma-Aldrich, V4505), 1:5000; anti-RPL26/RPL26L1 (Abcam, ab59567), 1:5000; anti-S6 ribosomal protein (Cell Signaling Technology (USA), 2217S), 1:5000; anti-RPL7A (Proteintech (USA), 15340-1-AP), 1:5000; anti-β-tubulin (DSHB (USA), E7), 1:5000; anti-V5 (Invitrogen (USA), R960-25), 1:3000; anti-NRF2 (Abcam, ab62352), 1:1000; anti-β-actin (Sigma-Aldrich, A5441-2ML), 1:5000; anti-ITPR2 (Alomone Labs (Israel), ACC-116), 1:1000. Membranes were washed with TBS containing 0.5% Tween-20 (TBST), incubated with secondary antibodies, and visualized using an enhanced chemiluminescence system. Protein levels were quantified by measuring band intensities using ImageJ software.

### Assessment of protein degradation and synthesis

When cells reached >90% confluency, the culture medium was replaced with serum-free MEM. After 72 hr under serum-free conditions, siRNA was transfected using Lipofectamine RNAiMAX (Thermo Fisher, 13778150) according to the manufacturer’s instructions.

To measure protein degradation, 48 hr after transfection, cells were treated with 10 μg/ml cycloheximide (CHX; Sigma-Aldrich, C1988). Cells were harvested at the indicated time points following CHX treatment and subjected to immunoblot analysis.

To assess protein synthesis, the SUnSET (surface sensing of translation) assay was performed based on the original method described by ^28^, with minor modifications. For *Tmem44* knockdown experiments, cells were treated with puromycin (Sigma-Aldrich, P8833) at 1 or 4 μg/ml for 30 min at 48 hr after transfection. For circadian translation experiments, cells were synchronized by serum shock as described above, and puromycin (4 μg/ml) was applied for 10 min prior to harvest at the indicated circadian time points. Cells were then harvested for immunoblotting. For detection of nascent peptides, anti-puromycin (12D10) (Sigma-Aldrich, MABE343), 1:3000, was used.

### *Tmem44* overexpression and immunocytochemistry

The pCMV-*Tmem44*-HA plasmid was custom-synthesized by Enzynomics (Republic of Korea) based on the pCMV-HA backbone (Clontech). Cells were seeded at a density of 1.2 × 10⁵ cells per well on 12-mm diameter poly-L-lysine–coated coverslips (Marienfeld (Germany), 0111520). When cells reached >90% confluency, the culture medium was replaced with serum-free MEM. After 72 hr under serum-free conditions, cells were transfected with pCMV-*Tmem44*-HA using Lipofectamine 2000 (Thermo Fisher, 11668019) according to the manufacturer’s instructions. Twelve hours after transfection, cells were fixed in 4% formaldehyde for 10 min at room temperature, washed twice with PBS, permeabilized in 1% PBST (1% Triton X-100 in PBS) for 5 min, and washed twice with PBS. Cells were then blocked with 1% bovine serum albumin (BSA) in 0.05% PBST for 30 min and incubated overnight at 4 °C with the following primary antibodies diluted in the same blocking buffer: anti-HA (3F10) (Roche (Switzerland), 12158167001), 1:200; anti-KDEL (10C3) (Enzo Biochem (USA), ADI-SPA-827), 1:100. After washing twice with 0.05% PBST, cells were incubated for 3 hr at room temperature with the following secondary antibodies diluted in 1% BSA in 0.05% PBST: goat anti-rat IgG–Alexa Fluor 488 (Thermo Fisher, A11006), 1:200; and goat anti-mouse IgG–Alexa Fluor 555 (Thermo Fisher, A21424), 1:100. Coverslips were washed twice with 0.05% PBST and mounted using DAPI-containing mounting medium (Abcam, ab104139).

Confocal images were acquired using an LSM 800 microscope (Carl Zeiss (Germany)) and processed with Zen software (Zen Digital Imaging for Light Microscopy, version 3.1; Carl Zeiss). Colocalization analysis was performed using the same software.

### Immunocytochemistry for TMEM44-miniTurbo localization and proximity labeling

The plasmids pEF1-EGFP-miniTurbo-V5 and pEF1-*Tmem44*-miniTurbo-V5 were custom-synthesized by VectorBuilder (USA). For immunocytochemistry, cells were seeded at a density of 2.5 × 10⁵ cells per well on 15-mm poly-L-lysine–coated coverslips (Marienfeld, 111550). When cells reached >90% confluency, the culture medium was replaced with serum-free MEM. After 72 hr under serum-free conditions, cells were transfected using Lipofectamine 2000 according to the manufacturer’s instructions. For localization analysis, cells were co-transfected with DsRed2-ER-5 (a gift from Michael Davidson, Addgene plasmid #55836 ^29^). For proximity labeling experiments, cells were transfected with pEF1-*Tmem44*-miniTurbo-V5 and treated with 100 µM biotin (Sigma-Aldrich, B4639) or DMSO for 1 hr at 48 hr post-transfection.

Cells were fixed with 4% formaldehyde for 10 min at room temperature, washed twice with PBS, and quenched with 100 mM glycine in PBS for 5 min. After additional PBS washes, cells were permeabilized with 0.05% saponin in PBS (PBSS) for 5 min and blocked with 5% BSA in 0.05% PBSS for 1 hr at room temperature. For localization analysis, cells were incubated overnight at 4 °C with anti-mCherry (Thermo Fisher, TFS-M11217), 1:300, and anti-V5 (Invitrogen, R960-25), 1:300, followed by incubation with goat anti-rat IgG–Alexa Fluor 594 (Thermo Fisher, A11007), 1:200, and goat anti-mouse IgG–Alexa Fluor 488 (Abcam, ab150113), 1:200. For proximity labeling, cells were incubated overnight at 4 °C with anti-V5 (Invitrogen, R960-25), 1:300, followed by incubation with goat anti-mouse IgG–Alexa Fluor 555 (Invitrogen, A21424), 1:200, and streptavidin–Alexa Fluor 488 (Thermo Fisher, S32354), 1:300, for 2 hr at room temperature. Coverslips were washed with 0.05% PBSS, rinsed with PBS, and mounted using DAPI-containing mounting medium (ab104139). Confocal images were acquired using an LSM 800 microscope (Carl Zeiss (Germany)) and processed with Zen software (Zen Digital Imaging for Light Microscopy, version 3.1; Carl Zeiss). Colocalization analysis was performed using the same software.

### Proximity labeling and streptavidin pulldown

Cells were seeded at a density of 2.5 × 10⁶ cells per 100-mm culture dish, and at least five dishes per condition were used to obtain sufficient protein for downstream analyses. Cells were cultured and transfected using Lipofectamine 2000 according to the manufacturer’s instructions. All procedures for TurboID-based proximity labeling and subsequent LC–MS/MS analysis were performed as previously described ^30^ with minor modification. Briefly, at 48 hr post-transfection, cells were treated with 100 µM biotin for 1 hr. Cells were then lysed in modified RIPA buffer (TurboID Mo-RIPA; 50 mM Tris-HCl pH 7.4, 150 mM NaCl, 0.1% SDS, 0.5% sodium deoxycholate, 1% Triton X-100) supplemented with protease inhibitor cocktail (Sigma-Aldrich, P8340) and 1 mM PMSF, followed by brief sonication.

For proteomics analysis, a total of 2250 µg of protein per condition was processed. Lysates were divided into three tubes (750 µg each), incubated with streptavidin-conjugated magnetic beads (Thermo Fisher, 88817) separately, and subsequently combined after pulldown. Streptavidin-conjugated magnetic beads were added at a ratio of 0.3 µl per µg protein for EGFP-miniTurbo-V5 samples and 0.2 µl per µg protein for TMEM44-miniTurbo-V5 samples, followed by incubation for 2 hr at 4 °C with end-over-end rotation. Biotinylated proteins were eluted by boiling the beads in 3x protein loading buffer supplemented with 2 mM biotin and 20 mM DTT at 95 °C for 10 min.

For quality control of proteomics samples, one fraction was separated on a 4–12% Tris-glycine gel (Thermo Fisher, XP04122BOX) and visualized using a silver staining kit (Thermo Fisher, 24612) to confirm protein distribution and concentration. In addition, streptavidin-enriched samples were analyzed by SDS-PAGE followed by immunoblotting for V5 and streptavidin to validate construct expression and biotinylation efficiency. 20 µg of the corresponding whole-cell lysates (input) were loaded alongside the pulldown samples.

For mass spectrometry, proteins were loaded onto a separate gel, and electrophoresis was stopped shortly after the samples entered the resolving gel. The gel was stained using a colloidal blue staining kit (Invitrogen, LC6025), and the stained lane was excised into approximately 1 cm segments. Gel pieces were submitted to Bertis Inc. (Republic of Korea) for in-gel digestion, peptide extraction, and subsequent LC–MS/MS analysis.

### LC–MS/MS analysis and data processing

In-gel digestion and LC–MS/MS analysis were performed by Bertis Inc. (Republic of Korea). Briefly, gel pieces were destained using 50% acetonitrile in 25 mM ammonium bicarbonate and dehydrated with 100% acetonitrile. Proteins were reduced with 10 mM dithiothreitol at 56 °C and alkylated with 25 mM iodoacetamide at room temperature in the dark. After dehydration, proteins were digested with trypsin (Promega) at 37 °C overnight. Peptides were extracted using acetonitrile containing 0.1% trifluoroacetic acid, dried under vacuum, and reconstituted in 0.1% trifluoroacetic acid.

Peptide samples were analyzed using an Ultimate 3000 UPLC system coupled to a FAIMS Pro Orbitrap Exploris 480 mass spectrometer (Thermo Fisher Scientific). Peptides were first loaded onto a trap column (Acclaim PepMap 100, 100 μm × 2 cm) and separated on an analytical column (EASY-Spray, 75 μm × 50 cm) at 50 °C with a flow rate of 0.3 μL/min. The mobile phases consisted of 0.1% formic acid and 5% dimethyl sulfoxide in water (buffer A) and 0.1% formic acid and 5% dimethyl sulfoxide in 80% acetonitrile (buffer B). Peptides were separated using a linear gradient of buffer B from 2% to 54% over 98 min, followed by a high-organic wash and re-equilibration.

Mass spectrometry was performed in data-dependent acquisition mode. MS1 scans were acquired at a resolution of 60,000 over a mass range of 350–1,800 m/z. MS2 spectra were acquired with an isolation window of 1.3 m/z, normalized collision energy of 28%, resolution of 15,000, an AGC target of 100%, and a maximum injection time of 22 ms.

Raw data were processed using the SAGE search engine (v0.14.7) against the UniProt Mus musculus database (UP000000589). Fully tryptic peptides with up to two missed cleavages were allowed. Carbamidomethylation of cysteine was set as a fixed modification, and oxidation of methionine was set as a variable modification. The false discovery rate was controlled at 1% at the spectrum, peptide, and protein levels. Differentially expressed proteins were defined using a permutation-based Student’s t-test (p < 0.05) and log2-median ratio thresholds based on the empirical distribution.

### Co-immunoprecipitation

Astrocytes were seeded at a density of 2.5 × 10⁶ cells per 100-mm culture dish. When cells reached >90% confluency, the medium was replaced with serum-free MEM, and after 72 hr under serum-free conditions, siRNA was transfected using Lipofectamine RNAiMAX according to the manufacturer’s instructions. Cells were harvested 72 hr after transfection and lysed in TurboID Mo-RIPA buffer, followed by brief sonication.

For immunoprecipitation, 500 μg of total protein was incubated with 6 μl of anti-TMEM44 antibody (this study, #11) overnight at 4 °C with end-over-end rotation. Protein A magnetic beads (Thermo Fisher, 88845; 50 μl) were then added and incubated for 3 hr at 4 °C with end-over-end rotation. Beads were washed three times with lysis buffer, and immune complexes were eluted by boiling in SDS-PAGE sample buffer and subjected to immunoblot analysis.

For immunoblot detection of immunoprecipitated samples, Rabbit TrueBlot® anti-rabbit IgG HRP (Rockland (USA), RK18-8816-31) was used for targets detected with rabbit primary antibodies that overlap with IgG heavy/light chains. For all other targets, HRP-conjugated secondary antibodies were used according to the host species of the primary antibodies (goat anti-rabbit IgG (Thermo Fisher, G-21234); goat anti-mouse IgG (Thermo Fisher, G-21040)).

### Statistics

Statistics were performed using GraphPad Prism 10software. Data are presented as mean ± SEM. Normality was assessed prior to statistical testing using the Shapiro–Wilk test. For normally distributed data, differences between two groups were analyzed using an unpaired two-tailed Student’s t-test, and comparisons among more than two groups were performed using one-way ANOVA followed by Tukey’s post hoc test. For experiments involving multiple conditions and time points, two-way ANOVA was used, followed by multiple comparisons tests as indicated in the figure legends. For data that did not meet normality assumptions, nonparametric tests were applied as appropriate. A *p* value < 0.05 was considered statistically significant.

## Results

### Identification of *Tmem44* as an astrocyte-enriched circadian transcript

We previously identified 412 oscillating transcripts in mouse cultured cortical astrocytes ^7^. Rhythmically expressed genes in astrocytes may not simply represent downstream outputs of the circadian clock but could themselves modulate TTFL in a cell-type specific manner ^8,31–33^. To identify such candidates, we narrowed this dataset by applying three independent criteria: mean expression exceeding TPM 10, high expression in adult mouse cortical astrocytes ^22^, and astrocyte-enriched expression relative to other cortical cell types ^23^. This stepwise filtering strategy identified 38 astrocyte-enriched circadian transcripts (Figure 1A, Table S1). From this set, we prioritized *Tmem44* for further analysis due to its strong circadian rhythmicity and limited functional characterization. *Tmem44* mRNA exhibited rhythmic oscillation with a phase similar to *Bmal1* and antiphasic to *Per2* and was upregulated with a flattened rhythm in the absence of *Bmal1* (Figure 1B).

**Figure 1.**
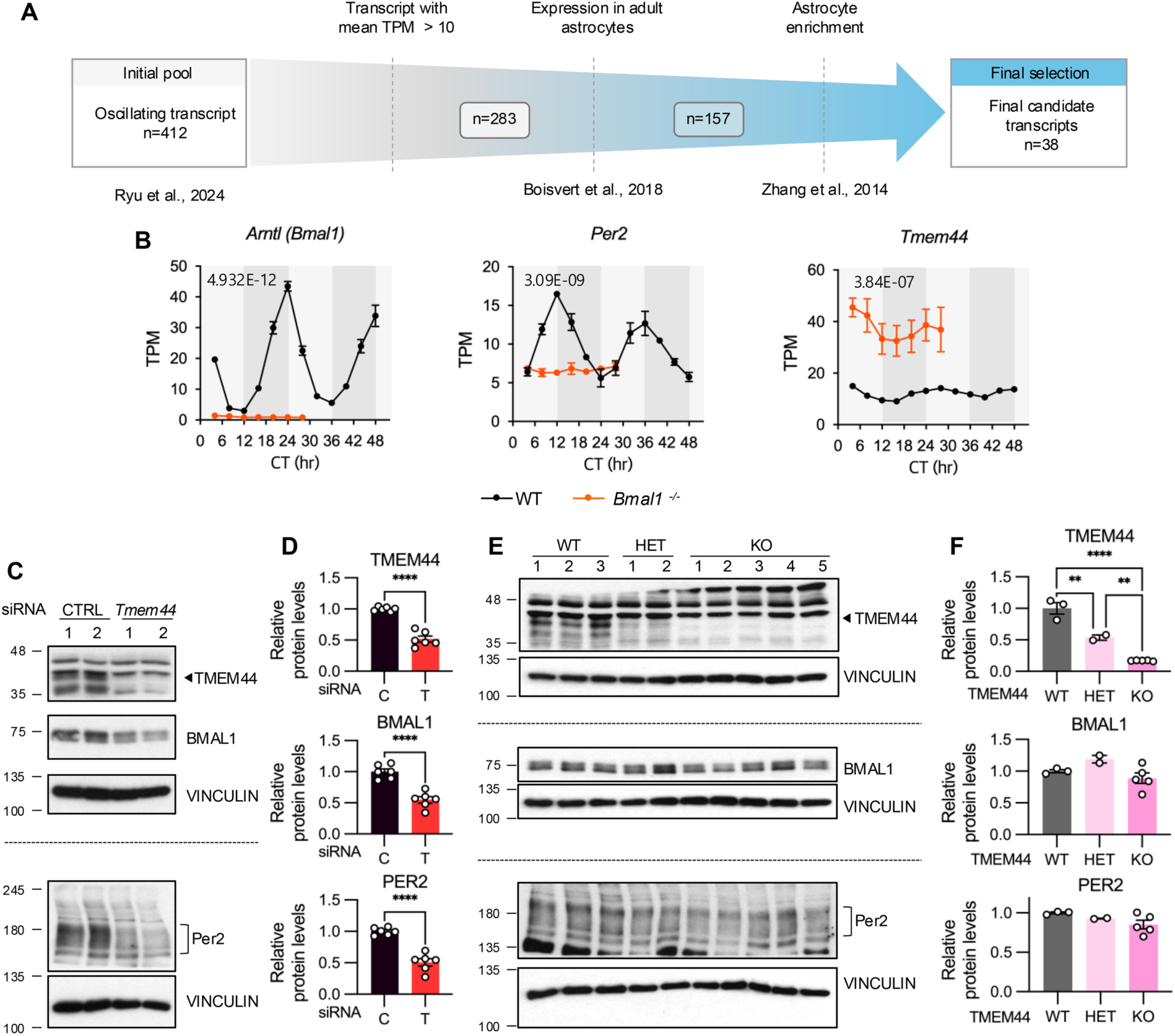
Identification of *Tmem44* as an astrocyte-enriched circadian transcript and its effect on clock protein abundance. (A) Schematic overview of the filtering strategy used to identify astrocyte-enriched circadian candidate transcripts. (B) Circadian expression profiles of *Bmal1*, *Per2*, and *Tmem44* in circadian-synchronized mouse cortical astrocytes ^7^. Data are presented as mean ± SEM. Meta2d p values are indicated within each plot. Light gray and dark gray backgrounds denote subjective day and subjective night, respectively. (C, D) Astrocytes were analyzed 48 h after transfection with control or *Tmem44*-targeting siRNA. (C) Representative western blot images showing BMAL1 and PER2 protein levels following siRNA-mediated knockdown of *Tmem44* in astrocytes. (D) Densitometric analysis of protein levels from (C). Protein levels were normalized to VINCULIN and further normalized to control astrocytes (set to 1). Control siRNA–treated astrocytes and *Tmem44* siRNA–treated astrocytes are labeled as C and T, respectively. Data are presented as mean ± SEM (n = 6). Statistical significance was determined by Student’s t-test. (E) Representative western blot images showing TMEM44, BMAL1, and PER2 protein levels in astrocytes derived from *Tmem44* wild-type, heterozygous, and knockout mice. (F) Densitometric analysis of protein levels from (E). Protein levels were normalized to VINCULIN and further normalized to wild-type astrocytes (set to 1). Data are presented as mean ± SEM (WT n = 3, HET n = 2, KO n = 5).

To assess whether *Tmem44* regulates the circadian clock, we examined core clock protein abundance upon *Tmem44* knockdown. *Tmem44* knockdown significantly reduced BMAL1 and PER2 protein levels, indicating that TMEM44 is required for normal clock protein abundance in astrocytes (Figure 1C and 1D). We next examined whether this effect was recapitulated in a genetic model by analyzing astrocytes derived from *Tmem44* wild-type, heterozygous, and knockout animals. TMEM44 protein levels decreased in a gene dosage–dependent manner; however, BMAL1 and PER2 protein levels were not altered (Figure 1E and 1F).

These results suggest that developmental or long-term compensatory mechanisms may mask the effect of constitutive *Tmem44* loss on clock protein abundance, whereas acute *Tmem44* knockdown reveals a requirement for TMEM44 in maintaining BMAL1 and PER2 levels Therefore, subsequent experiments were performed using *Tmem44* knockdown astrocytes.

### *Tmem44* knockdown reduces BMAL1 and PER2 protein abundance and dampens transcriptional rhythms

To examine how *Tmem44* knockdown affects circadian dynamics across circadian cycle, we analyzed BMAL1 and PER2 in synchronized astrocytes. In control astrocytes, BMAL1 mobility shifts indicative of phosphorylation and PER2 protein levels displayed robust circadian variation ^34,35^. In *Tmem44* knockdown astrocytes, overall BMAL1 and PER2 protein levels were significantly reduced across all circadian time points; however, the temporal phosphorylation patterns of both proteins were preserved (Figure 2A and 2B). We next examined whether this reduction in protein abundance was due to altered transcription. Quantitative PCR analysis revealed that *Tmem44* knockdown did not significantly affect *Bmal1* or *Per2* mRNA levels (Figure 2C). However, real-time bioluminescence recording of a *Per2*-luciferase reporter, which provides higher temporal resolution than endpoint qPCR, showed that *Tmem44* knockdown significantly reduced oscillation amplitude and advanced the circadian phase without altering the period (Figure 2D and 2E). Together, *Tmem44* knockdown reduces clock protein abundance without altering steady-state mRNA levels, and the resulting reduction likely feeds back onto the TTFL to dampen transcriptional rhythms.

**Figure 2.**
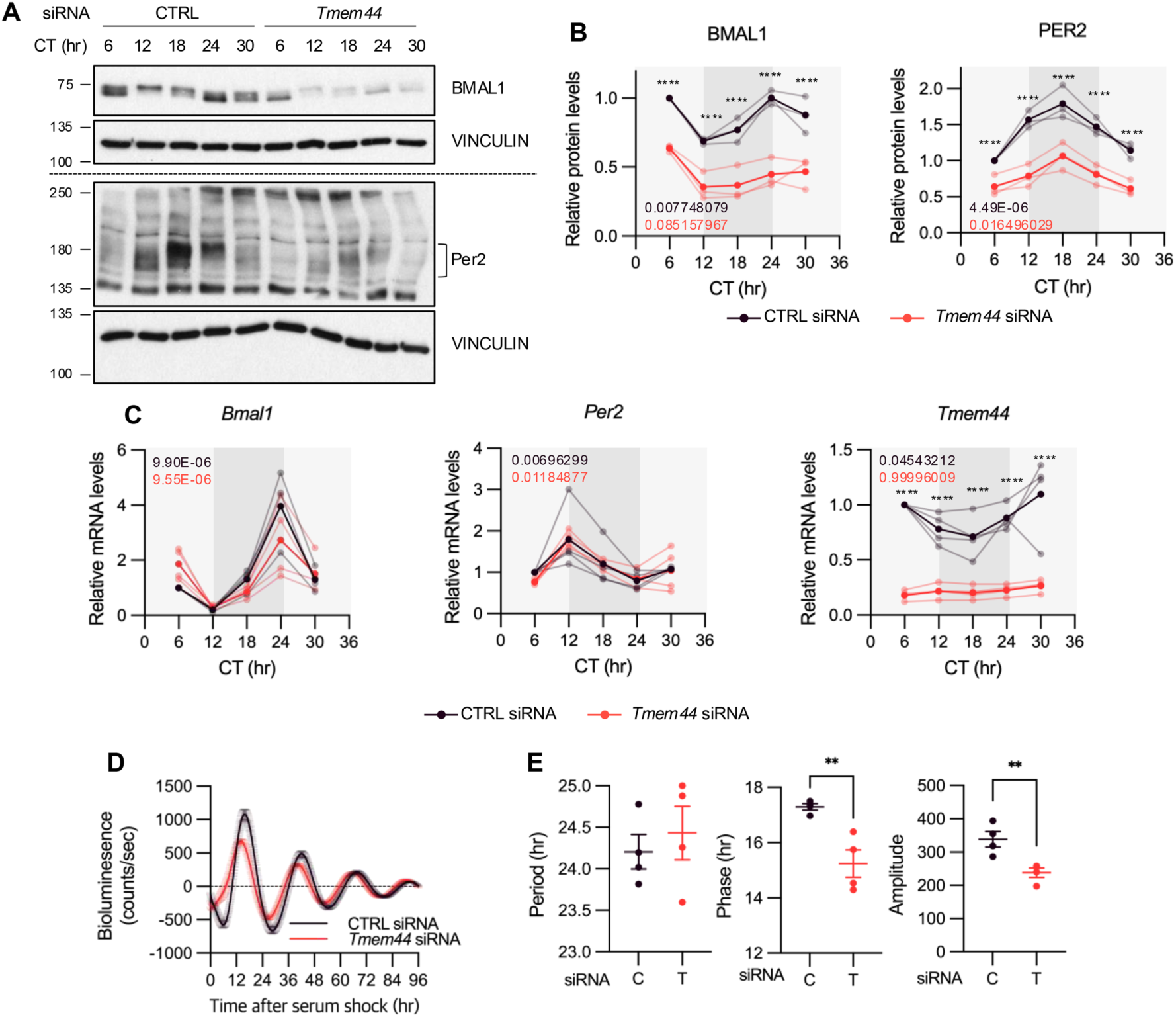
*Tmem44* knockdown reduces BMAL1 and PER2 protein abundance without altering their mRNA levels. Astrocytes were transfected with control siRNA or *Tmem44*-targeting siRNA and synchronized by serum shock 48 h after transfection. Cells were harvested at the indicated circadian time points for analysis. Control astrocytes and *Tmem44* knockdown astrocytes are shown in dark purple and red, respectively. (A) Representative western blot images showing BMAL1 and PER2 protein levels across circadian time. (B) Densitometric analysis of protein levels from (A). Protein levels were normalized to VINCULIN, and values were further normalized to the CT6 time point of control astrocytes (set to 1). Data are presented as mean ± SEM (n = 4). Meta2d p values are indicated within the graph. Light gray and dark gray backgrounds denote subjective day and subjective night, respectively. Statistical significance was determined by two-way ANOVA, and asterisks indicate results from multiple comparisons (****p < 0.0001). (C) Circadian mRNA expression profiles of *Bmal1* and Per2. Data are presented as mean ± SEM (n = 4). Values were normalized to the CT6 time point of control astrocytes (set to 1). Meta2d p values are indicated within each graph. Light gray and dark gray backgrounds denote subjective day and subjective night, respectively. Statistical significance was determined by two-way ANOVA, and asterisks indicate results from multiple comparisons (****p < 0.0001). (D) Baseline-subtracted luminescence recordings showing circadian oscillations from Per2-luciferase reporter astrocytes Data are presented as mean ± SEM (n = 4). (E) Analysis of circadian parameters derived from (D), including period, phase, and amplitude. Control siRNA–treated astrocytes and *Tmem44* siRNA–treated astrocytes are labeled as C and T, respectively. Data are presented as mean ± SEM (n = 4). Statistical significance was determined by two-way ANOVA, and asterisks indicate results from multiple comparisons (**p < 0.01).

### *Tmem44* protein is constitutively expressed and localizes to the endoplasmic reticulum

*Tmem44* encodes a predicted multi-pass membrane protein with seven transmembrane helices, as supported by independent topology prediction algorithms (DeepTMHMM and Phobius) ^36,37^. Despite this predicted membrane topology, the biological function of *Tmem44* remains largely uncharacterized. We therefore examined its protein expression and subcellular localization in astrocytes. Although *Tmem44* mRNA oscillates robustly under circadian clock control (Figure 1B and 2C), TMEM44 protein levels remained constant across circadian time points in both control and *Tmem44* knockdown astrocytes (Figure 3A and 3B). In *Bmal1*⁻/⁻ astrocyte cultures, where *Tmem44* mRNA was elevated and arrhythmic (Figure 1B), TMEM44 protein abundance was correspondingly increased compared to wild-type controls (Figure 3C and 3D). These results indicate that TMEM44 protein abundance reflects steady-state *Tmem44* mRNA levels but does not exhibit circadian oscillation, suggesting that clock-dependent regulation of *Tmem44* occurs primarily at the transcript level.

**Figure 3.**
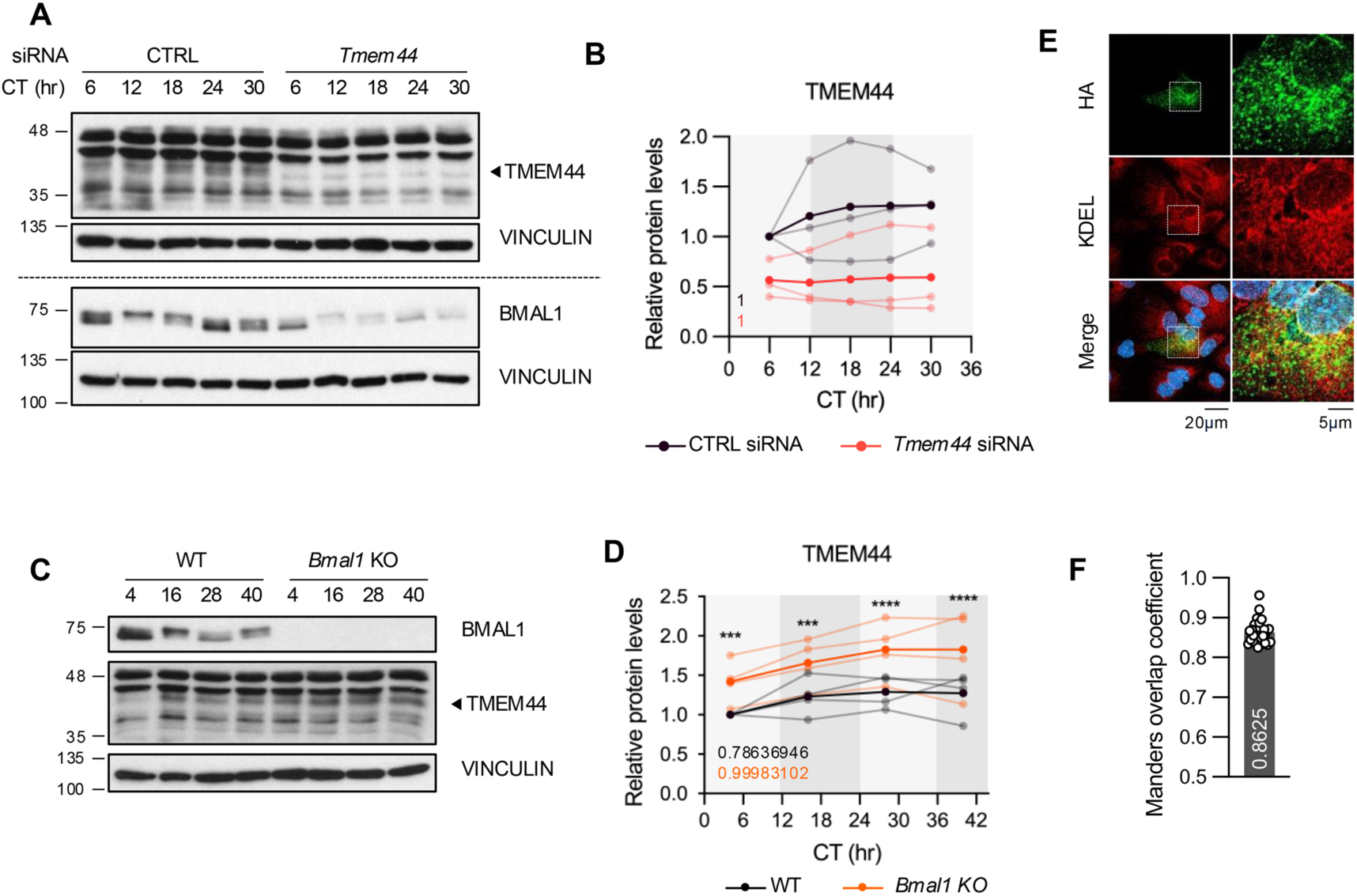
TMEM44 protein is constitutively expressed and localizes to the ER in astrocytes. (A) Representative western blot images showing TMEM44 and BMAL1 protein levels across circadian time in serum shock–synchronized control and *Tmem44* knockdown astrocytes. (B) Densitometric analysis of TMEM44 protein levels from (A). Protein levels were normalized to VINCULIN and further normalized to the CT4 time point of control astrocytes (set to 1). Data are presented as mean ± SEM (n = 3). Meta2d p values are indicated within the graph. Light gray and dark gray backgrounds denote subjective day and subjective night, respectively. Statistical significance was determined by two-way ANOVA. (C) Representative western blot images showing TMEM44 protein levels in serum shock-synchronized astrocytes derived from WT and *Bmal1*⁻/⁻ mice at selected time points. (D) Densitometric analysis of TMEM44 protein levels from (C). Protein levels were normalized to VINCULIN and further normalized to the CT4 time point of WT astrocytes (set to 1). WT and *Bmal1*⁻/⁻ astrocytes are shown in black and orange, respectively. Data are presented as mean ± SEM (n = 4). Meta2d p values are indicated within the graph. Statistical significance was determined by two-way ANOVA, and asterisks indicate results from multiple comparisons (*p < 0.05, **p < 0.01, ***p < 0.001, ****p < 0.0001). (E) Representative immunocytochemistry images of astrocytes transfected with HA-tagged TMEM44 (TMEM44–HA). Cells were fixed 12 h after transfection. (F) Colocalization analysis between TMEM44-HA and KDEL signals shown in (E). The Manders overlap coefficient was calculated from the images (n = 30).

To determine the subcellular localization of TMEM44 in astrocytes, we performed immunocytochemical analysis using HA-tagged TMEM44, as the antibody was not suitable for immunofluorescence. TMEM44-HA showed predominant colocalization with the ER marker KDEL (Figure 3E), with a Manders overlap coefficient of 0.86 (Figure 3F), indicating that TMEM44 is an ER-resident protein.

### *Tmem44* regulates clock protein abundance through protein synthesis

To investigate how *Tmem44* knockdown reduces clock protein abundance, we assessed the contribution of protein degradation and protein synthesis. Cycloheximide (CHX) chase experiments revealed comparable degradation kinetics of BMAL1, and PER2 between control and *Tmem44* knockdown astrocytes, despite lower baseline protein levels in the knockdown condition (Figure 4A and 4B), indicating that *Tmem44* knockdown does not affect clock protein degradation.

**Figure 4.**
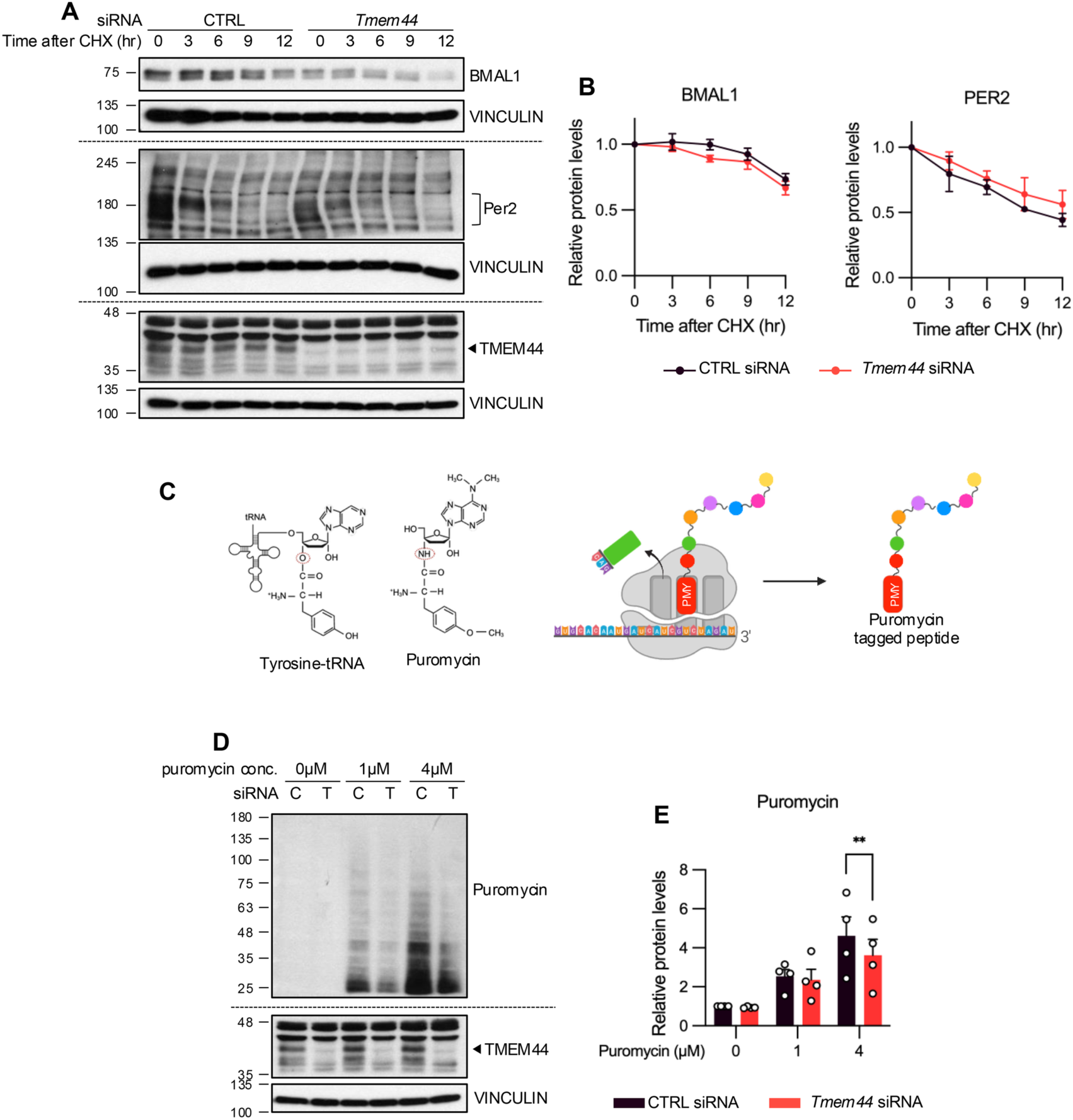
TMEM44 regulates clock protein abundance by supporting protein synthesis rather than protein stability in astrocytes. Astrocytes were analyzed 48 h after transfection with control or *Tmem44*-targeting siRNA. (A, B) Cycloheximide (CHX) chase assay to assess protein stability. Cells were treated with CHX (10 μg/mL) for the indicated durations. (A) Representative western blot images. (B) Densitometric analysis of protein levels from (A). Protein levels were normalized to VINCULIN, and values were further normalized to the 0 h time point of each condition (set to 1). Data are presented as mean ± SEM (n = 3). (C–E) SUnSET assay to assess global protein synthesis in control astrocytes and *Tmem44* knockdown astrocytes. (C) Schematic diagram illustrating the principle of the SUnSET assay. (D) Representative western blot images. Control siRNA–treated astrocytes and *Tmem44* siRNA–treated astrocytes are labeled as C and T, respectively. (E) Densitometric analysis of puromycin incorporation from (D). Puromycin signals were normalized to VINCULIN, and values were further normalized to the control astrocytes treated with 0 μM puromycin (set to 1). Data are presented as mean ± SEM (n = 4). Statistical significance was determined by two-way ANOVA, and asterisks indicate results from multiple comparisons (**p < 0.01).

We therefore examined global protein synthesis using the surface sensing of translation (SUnSET) assay, which measures nascent translation via puromycin incorporation into elongating polypeptides (Figure 4C) ^28^. Global translation was significantly reduced in *Tmem44* knockdown astrocytes compared to controls (Figure 4D and 4E).

Given the reduction in global translation, we next asked whether this translational suppression extends beyond clock proteins by examining a select panel of additional proteins. ITPR2 and NRF2 were reduced in *Tmem44* knockdown astrocytes, whereas cytoskeletal proteins (Vinculin, Tubulin, and Actin) and CREB were unaffected (Supplementary Figure 2A and 2B) indicating the effect of *Tmem44* knockdown on protein abundance extends beyond clock proteins yet does not uniformly affect all proteins.

Collectively, these data demonstrate that *Tmem44* knockdown reduces clock protein abundance by impairing protein synthesis.

### Proximity labeling identifies TMEM44-associated translational machinery at the ER

To investigate the molecular basis of TMEM44-dependent regulation of protein synthesis, we performed miniTurbo-based proximity labeling to identify proteins in the vicinity of TMEM44 in astrocytes (Figure 5A) ^38^. A TMEM44–miniTurbo–V5 fusion construct was expressed in astrocyte cultures, with EGFP–miniTurbo–V5 serving as a cytosolic control. TMEM44–miniTurbo–V5 localized predominantly to the ER (Manders coefficient = 0.88), consistent with TMEM44-HA (Figure 3E and 3F), while EGFP–miniTurbo–V5 showed a diffuse cytoplasmic distribution (Figure 5B and 5C), indicating that the fusion to miniTurbo preserved the native localization of TMEM44. Treatment with 100 µM biotin for 60 min yielded robust, spatially restricted biotinylation (Supplementary Figure 3A – 3D), and streptavidin pulldown recovered biotinylated proteins only in biotin-treated samples (Supplementary Figure 3E and Figure 5D). These samples were analyzed by LC–MS/MS.

**Figure 5.**
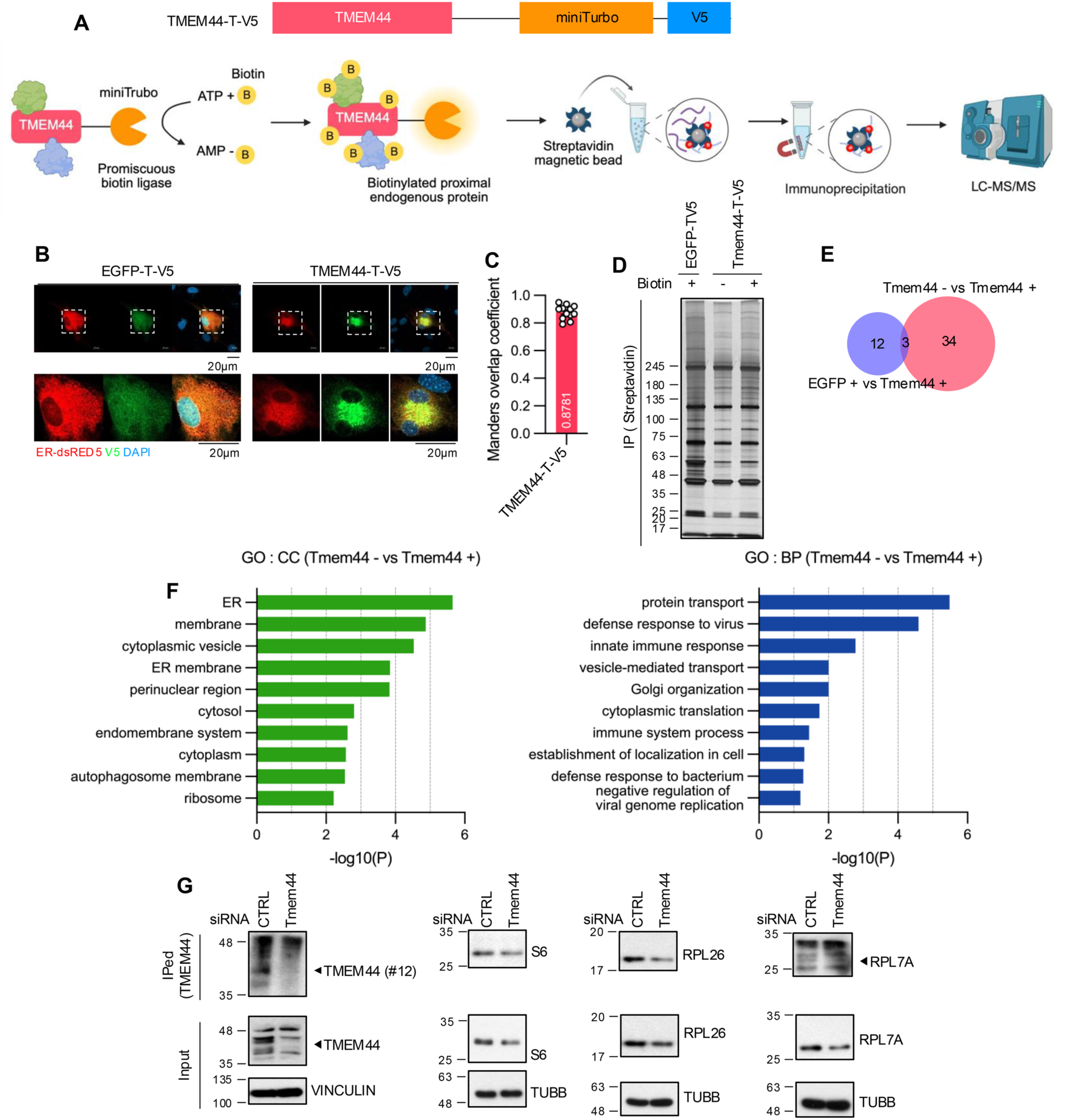
Proximity labeling identifies TMEM44-associated proteins in astrocytes. (A–D) *TMEM44*–miniTurbo–V5 and EGFP–miniTurbo–V5 are abbreviated as *TMEM44*–T–V5 and EGFP–T–V5, respectively, in the figure. (A) Schematic diagram of *TMEM44*–miniTurbo–V5–based proximity labeling and downstream workflow. (B) Representative immunocytochemistry images showing subcellular localization of EGFP–miniTurbo–V5 and TMEM44–miniTurbo–V5. (C) Manders overlap coefficient analysis of TMEM44–miniTurbo–V5 colocalization (n = 12). (D) Silver staining of streptavidin pull-down samples following biotin labeling. (E) Venn diagram showing differentially enriched proteins identified by proteomic analysis. (F) Gene ontology (GO) analysis of proteins enriched in the TMEM44+ condition compared to TMEM44−. Selected GO terms were abbreviated in the figure, including “perinuclear region of cytoplasm” as “perinuclear region” and “endoplasmic reticulum” as “ER.” (G) Immunoprecipitation of TMEM44 using a TMEM44 (#11) antibody, followed by immunoblotting for ribosomal proteins (RPS6, RPL26, and RPL7A) and TMEM44 in astrocytes. Input and immunoprecipitated (IP) samples are shown.

We compared three conditions: biotin-treated EGFP–miniTurbo–V5 (EGFP+), untreated TMEM44–miniTurbo–V5 (TMEM44−), and biotin-treated TMEM44–miniTurbo–V5 (TMEM44+). This yielded 15 proteins enriched in the EGFP+ vs TMEM44+ comparison and 37 proteins in the TMEM44− vs TMEM44+ comparison, with minimal overlap between datasets (Figure 5E). EGFP–miniTurbo–V5 showed higher expression and stronger global biotinylation than TMEM44-miniTurbo-V5, likely reflecting increased background (Supplementary Figure 3A and 3B). Subsequent analyses therefore focused on the TMEM44− vs TMEM44+ dataset as a more reliable source of TMEM44-proximal candidates (Table S2).

Among the 37 proteins identified, 34 were enriched in the TMEM44+ condition, including ribosomal proteins (Rpl7a, Rpl26, and Rps6) and ER-associated translation machinery components (Ckap4, Rpn2, Tram1). Gene Ontology (GO) analysis using the DAVID platform ^39,40^ revealed strong enrichment of ER and ribosome-related terms In the Cellular Component category, and cytoplasmic translation in the Biological Process category (Figure 5F), consistent with a role for TMEM44 in ER-associated translation. In addition, terms related to protein transport, antiviral response, and innate immunity were enriched in the Biological Process category (Figure 5F).

To validate the physical association between TMEM44 and ribosomal proteins identified by proximity labeling, we performed co-immunoprecipitation in control and *Tmem44* knockdown astrocytes. RPS6, RPL26, and RPL7A were co-immunoprecipitated with TMEM44 in control astrocytes, and their recovery was reduced in *Tmem44* knockdown astrocytes, consistent with the reduction in TMEM44 bait levels (Figure 5G).

Collectively, these results demonstrate that TMEM44 is physically associated with ribosomal proteins at the ER, positioning it to regulate protein synthesis in astrocytes.

### Circadian regulation of protein translation in astrocytes requires TMEM44

Given that TMEM44 associates with ribosomal proteins and its knockdown regulates global protein synthesis, we asked whether TMEM44 is required for circadian regulation of translation. Previous reports have demonstrated that global translation exhibits circadian oscillation in liver, brain, SH-SY5Y and U2OS ^13–15^, and we reasoned that TMEM44 at the ER may contribute to such rhythmic translational output in astrocytes.

To test this, we performed the SUnSET assay across circadian time points in synchronized astrocytes. In control astrocytes, BMAL1 phosphorylation confirmed robust circadian oscillation, and puromycin incorporation displayed clear circadian rhythmicity, peaking during the subjective day and reaching a trough during the subjective night (Figure 6A and 6B). In *Tmem44* knockdown astrocytes, puromycin incorporation was not only reduced in overall levels, consistent with our earlier findings (Figure 4D) but also lost its circadian oscillation, resulting in a flattened temporal profile (Figure 6A and 6B).

**Figure 6.**
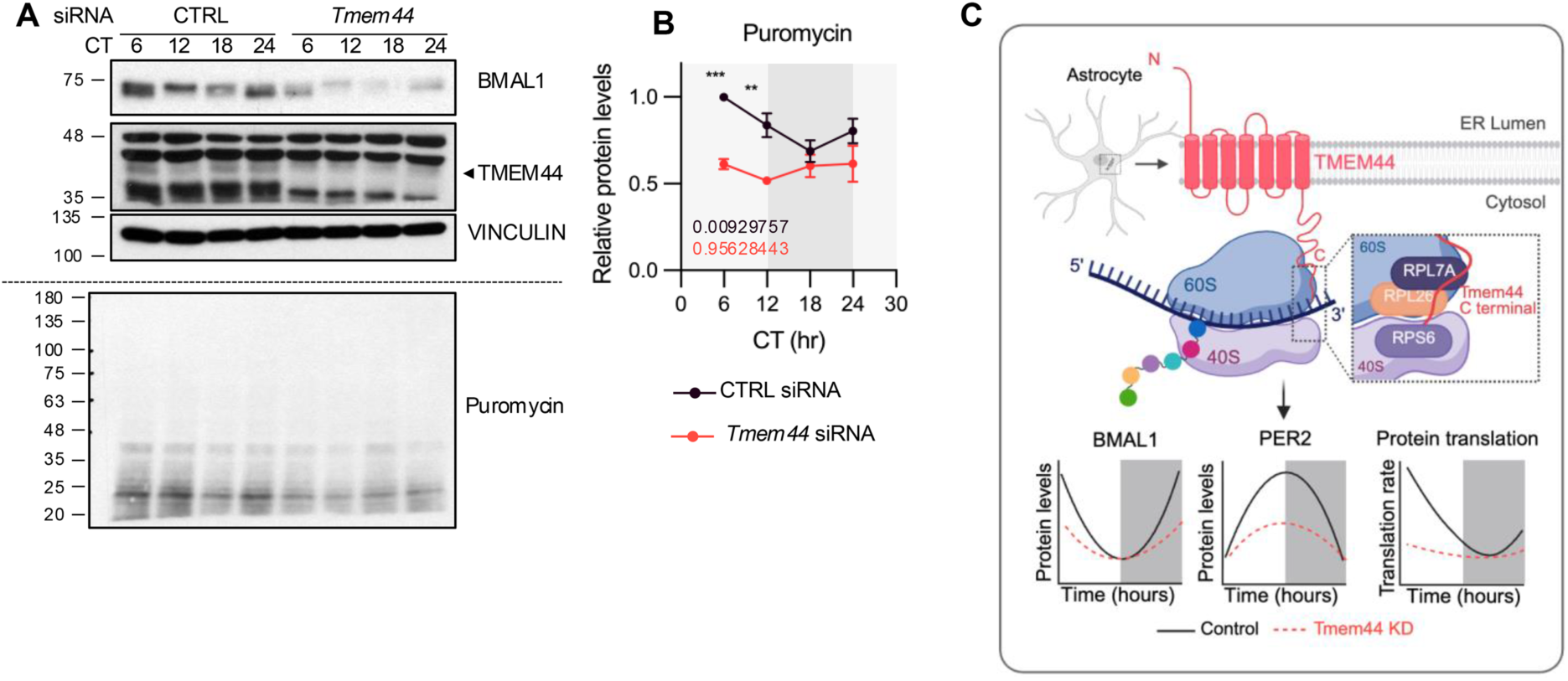
*Tmem44* is required for circadian regulation of protein synthesis in astrocytes. Astrocytes were analyzed 48 h after transfection with control or *Tmem44*-targeting siRNA, synchronized by serum shock, and subjected to SUnSET assays at the indicated circadian times (CT6, CT12, CT18, and CT24) following puromycin treatment (4 μM, 10 min). (A) Representative western blot images showing puromycin incorporation and protein levels. (B) Densitometric analysis of BMAL1, TMEM44, and puromycin signals from (A). All signals were normalized to VINCULIN, and values were further normalized to the CT6 time point of control astrocytes (set to 1). Data are presented as mean ± SEM (n = 4). Meta2d p values are indicated within the graph. Light gray and dark gray backgrounds denote subjective day and subjective night, respectively. Statistical significance was determined by two-way ANOVA, and asterisks indicate results from multiple comparisons. (C) Schematic model summarizing the role of TMEM44 in astrocyte circadian protein synthesis. The ER-resident protein TMEM44 associates with ribosomal proteins, including RPS6, RPL26, and RPL7A and supports circadian oscillation of global protein translation. *Tmem44* knockdown abolishes translational rhythmicity and reduces BMAL1 and PER2 protein levels.

These results demonstrate that *Tmem44* is required for circadian regulation of protein synthesis in astrocytes and further suggest that rhythmic translational activity mediated by TMEM44 is necessary for sustaining high-amplitude oscillations of clock proteins (Figure 6C).

## Discussion

The circadian clock coordinates cellular physiology through rhythmic transcriptional programs in a highly cell type–specific manner ^41,42^. Although circadian transcription has been extensively characterized across tissues, considerably less is known about how circadian clocks regulate protein synthesis, particularly in astrocytes. Here, through functional screening of astrocyte-enriched circadian candidates, we identify TMEM44—a previously uncharacterized ER—resident membrane protein-as a regulator of astrocyte circadian clock function. TMEM44 is required to sustain BMAL1 and PER2 protein levels and to maintain high-amplitude *Per2*-luciferase rhythms, establishing it as a regulator of clock protein abundance and circadian rhythm amplitude in astrocytes.

*Tmem44* mRNA exhibits robust circadian oscillation with a phase similar to *Bmal1*, and this rhythm is abolished in *Bmal1*-deficient astrocytes, which instead display an elevated, non-oscillating baseline. These features are consistent with indirect BMAL1-dependent repression of *Tmem44*, potentially through the REV-ERB/ROR axis. In this axis, BMAL1 drives expression of REV-ERB repressors, which in turn suppress target genes containing REV-ERB/ROR response elements (RREs) ^2,43,44^.Consistent with this possibility, *in silico* inspection of the *Tmem44* upstream region identified two putative RRE motifs. However, direct identification of the *cis*-regulatory elements driving *Tmem44* rhythmicity will require promoter-level analysis in future studies. Although *Tmem44* is a clock-controlled output, it feeds back to support clock function. Knockdown of *Tmem44* reduced BMAL1 and PER2 protein levels and produced a low-amplitude, phase-advanced *Per2*-luc rhythm. Mechanistically, reduced BMAL1 would be expected to limit transcriptional drive on *Per* promoters, thereby lowering oscillation amplitude ^2,45,46^, while reduced PER2 may accelerate the termination of the negative feedback cycle, thereby advancing circadian phase ^47,48^. Given that *Tmem44* is enriched in the adult astrocytes, these findings identify an astrocyte-enriched, clock-controlled component that in turn sustains high-amplitude oscillation of core clock proteins.

Multiple lines of evidence converge on a translational role for TMEM44. First, *tmem44* knockdown reduced BMAL1 and PER2 abundance, and this effect extended beyond clock proteins (Supplementary figure 2). Second, SUnSET analysis demonstrated that *Tmem44* knockdown reduced global nascent peptide synthesis. Third, miniTurbo proximity labeling and co-IP analysis recovered ribosomal subunits including RPS6, RPL26, and RPL7A, indicating that TMEM44 physically associates with the translational machinery. Together, these findings identify TMEM44 as a component of the ER-associated translation apparatus. The localization of TMEM44 to the ER (Figure 3E and 3F) was initially unexpected given its effect on BMAL1 and PER2, which are neither membrane-bound nor secretory. However, it is now well established that ER-associated translation is not restricted to secretory and membrane proteins. Ribosome profiling and subcellular fractionation studies indicate that a broad range of mRNAs are detected in ER-associated ribosome fractions, suggesting that translation is distributed across multiple subcellular compartments rather than being confined to the cytosol ^49–51^. Consistent with this view, the published dataset Horste et al. ^51^ reports that transcripts encoding core clock proteins, including *Arntl* (*Bmal1*) and *Per2*, are detected in both cytosolic and ER-associated fractions at comparable levels, indicating that their translation is not exclusively cytosolic. Similar patterns are observed for additional TTFL-associated transcription factors such as *Dbp*, *Dec1*, and *Nfil3*, as well as for other transcription factor families including ATF, CREB3, and ZNF proteins, many of which exhibit detectable or even relatively enriched signals in ER-associated fractions. Together, these observations support a model in which translation of a wide range of non-secretory proteins, including transcriptional regulators, can occur in association with ER-bound ribosomes. Under this framework, the ER can function as a spatially distributed translational platform rather than a compartment dedicated solely to secretory protein synthesis. In this context, an ER-resident protein such as TMEM44 is well positioned to influence protein synthesis broadly, including that of non-secretory clock proteins.

The molecular features of TMEM44 further support a role in organizing translational machinery at the ER. The cytosolic C-terminus of TMEM44 contains intrinsically disordered segments, which often serve as flexible platforms for protein–protein interactions and regulatory signaling ^52,53^. Cytosolic tails of membrane proteins frequently harbor short linear motifs that mediate dynamic interactions with binding partners ^54,55^. In this context, the TMEM44 C-terminus may serve as an interaction interface that recruits or organizes translation-associated factors at ER subdomains. This model is consistent with our observation that TMEM44 associates with ribosomal proteins yet is unlikely to function as a structural component of the ribosome itself.

The effect of *Tmem44* knockdown on protein abundance was not uniform (Supplementary figure 2). Because SUnSET measures nascent protein synthesis over a short time window, proteins with rapid turnover and strong dependence on continuous translation-such as BMAL1, PER2, ITPR2, and NRF2-are expected to be more sensitive to a reduction in translation than proteins with slower turnover, whose steady-state levels are buffered by their longer half-lives. This differential response may account for the selective reductions we observed and indicates that TMEM44 acts as a facilitator of translation rather than on obligate core component of the ribosome.

The core circadian machinery is regulated not only transcriptionally but also through translational control. *Ataxin-2* has been shown to support robust circadian rhythms in *Drosophila* ^56,57^ and mammals ^15^. In SCN neurons, ATXN2 and ATXN2L form rhythmic biomolecular condensates that recruit ribosomes and *Per* mRNAs, thereby driving oscillatory PER translation^15^. Additional translational regulators, including LARK and hnRNP Q (SYNCRIP), have been implicated in the translational control of circadian clock components across species ^58–60^. Our finding that TMEM44, an astrocyte-enriched ER-resident integral membrane protein, is required to maintain BMAL1 and PER2 levels places it within these translational regulators of the clock. Notably, Zhuang et al. (2023) further reported that ATXN2/ATXN2L condensates are closely intertwined with the ER, indicating that condensate-based and ER membrane-based translational machineries are spatially integrated rather than operating as separate systems. The rhythmic, condensate-driven translational activation of *Per* mRNAs by ATXN2/ATXN2L and the ER membrane-localized translational control we describe for TMEM44 may therefore both contribute to rhythmic translation of clock proteins, acting at physically connected subcellular platforms rather than at independent sites. Because TMEM44 is enriched in astrocytes and expressed at low levels in neurons ^23^, this cooperative arrangement between ATXN2/ATXN2L condensates and ER membrane-localized TMEM44 may operate predominantly in astrocytes; in neurons and other cell types where TMEM44 is sparse, functionally analogous ER-resident proteins may substitute for TMEM44 to support condensate-coupled translational control of clock components. More broadly, cell-type-specific clock amplitude is shaped, in part, by the local complement of translational facilitators distributed across these interconnected compartments.

The circadian oscillation of protein synthesis observed in astrocytes (Figure 6B) suggests that translational control represents an additional layer of circadian regulation in glia. In mammals, periods associated with sleep or rest are often accompanied by elevated biosynthetic activity, including increased protein synthesis ^61–63^. The higher translational activity we observed during the subjective day in cultured astrocytes is consistent with a temporally coordinated anabolic program. Sleep is also associated with synaptic remodeling, including both synaptic weakening and strengthening ^64^, processes that depend in part on regulated protein synthesis. Astrocytes, which structurally and metabolically support synapses and contribute to synapse remodeling by providing proteins or modulating the extracellular environment in a time-of-day-dependent manner ^65,66^. Our results raise the possibility that TMEM44-dependent translation helps couple astrocyte protein synthesis to circadian metabolic states and synapse-related remodeling, thereby linking circadian transcriptional programs to downstream translational output in astrocytes. These interpretations, however, should be considered in light of an important limitation of our study. All experiments were performed in primary cortical astrocyte cultures, which lack the neuronal, vascular, and systemic inputs that shape astrocyte physiology *in vivo*. Future studies using astrocyte-specific *Tmem44* manipulation *in vivo* will be required to test whether TMEM44-dependent translation contributes to circadian regulation of astrocyte function, sleep-associated biosynthetic states, and synaptic remodeling in the intact brain. Despite these limitations, our identification of TMEM44 as an ER-resident, astrocyte-enriched translational regulator embedded within the circadian feedback architecture defines a new molecular entry point for dissecting how circadian clocks shape protein synthesis in glia.

## Acknowledgements

We are grateful to members of Eun Young Kim laboratory for helpful discussions and to Jaerack Chang and Eun Jeong Lee for insightful comments. This research was supported by a grant from the Korea Dementia Research Project through the Korea Dementia Research Center (KDRC), funded by the Ministry of Health and Welfare and the Ministry of Science and ICT, Republic of Korea (RS-2025-02215497) and a grant from the National Research Foundation of Korea (NRF), funded by the Ministry of Science and ICT, Republic of Korea (RS-2025-24534961, RS-2026-25485901) to E.Y.K., and supported by Basic Science Research Program through the National Research Foundation of Korea(NRF) funded by the Ministry of Education (RS-2025-25421460) to J.E.R

## Author contributions

J.E.R. and E.Y.K. designed the research; J.E.R., M.P., and H.W.R. performed the research; J.E.R., M.P., J.-H.L., and E.Y.K. analyzed the data; and J.E.R. and E.Y.K. wrote the manuscript.

## Data availability

The proteomics datasets generated during the current study will be deposited in the PRIDE repository prior to publication. Other data supporting the findings of this study are available from the corresponding author upon reasonable request.

## Supplementary Materials for

The astrocyte-enriched gene Tmem44 regulates circadian protein translation in mouse astrocytes

**Table S1.**
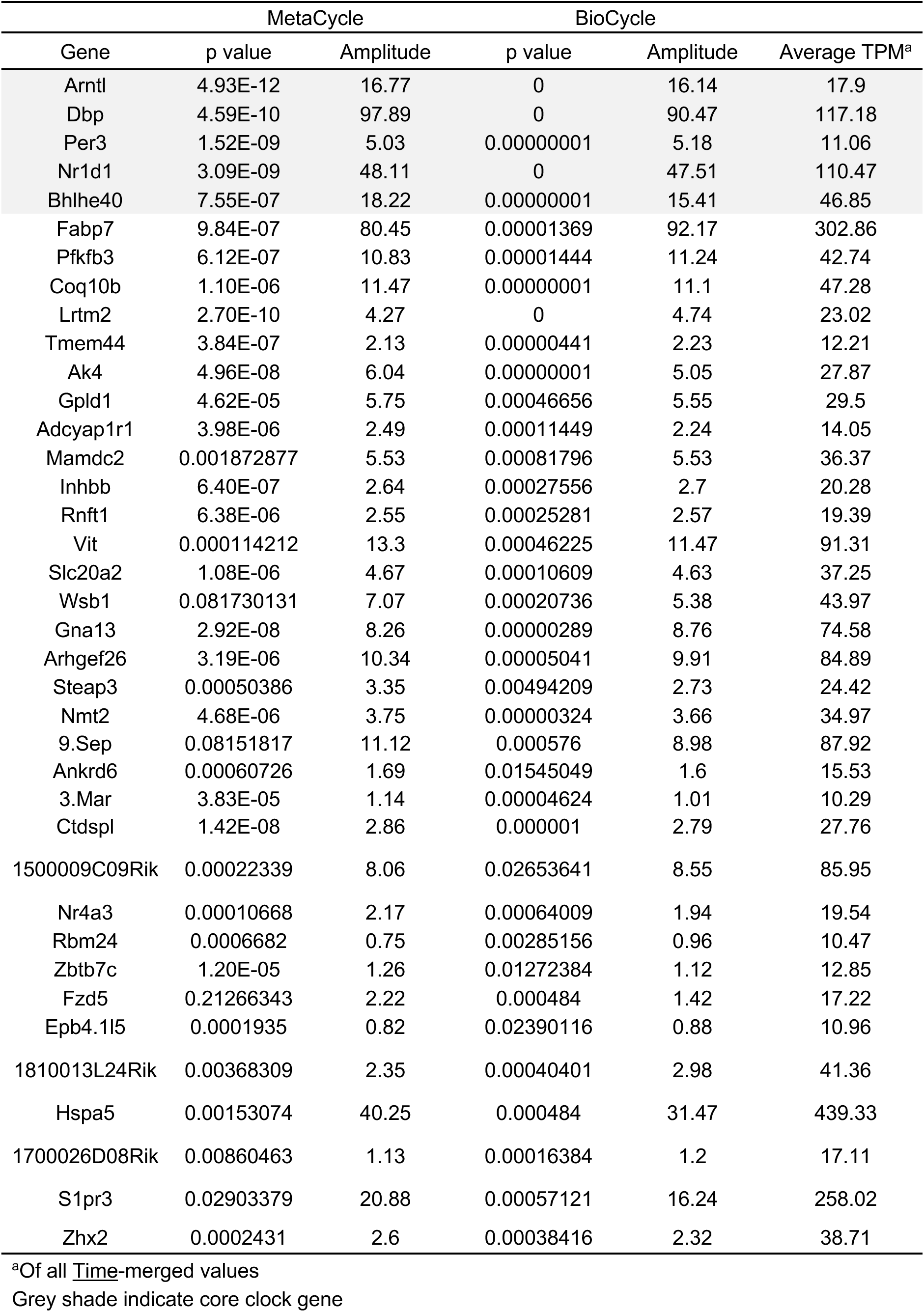
Astrocyte-enriched circadian transcripts. List of the 38 astrocyte-enriched circadian transcripts, ranked by MetaCycle p value. Grey shading indicates core clock genes

**Table S2.**
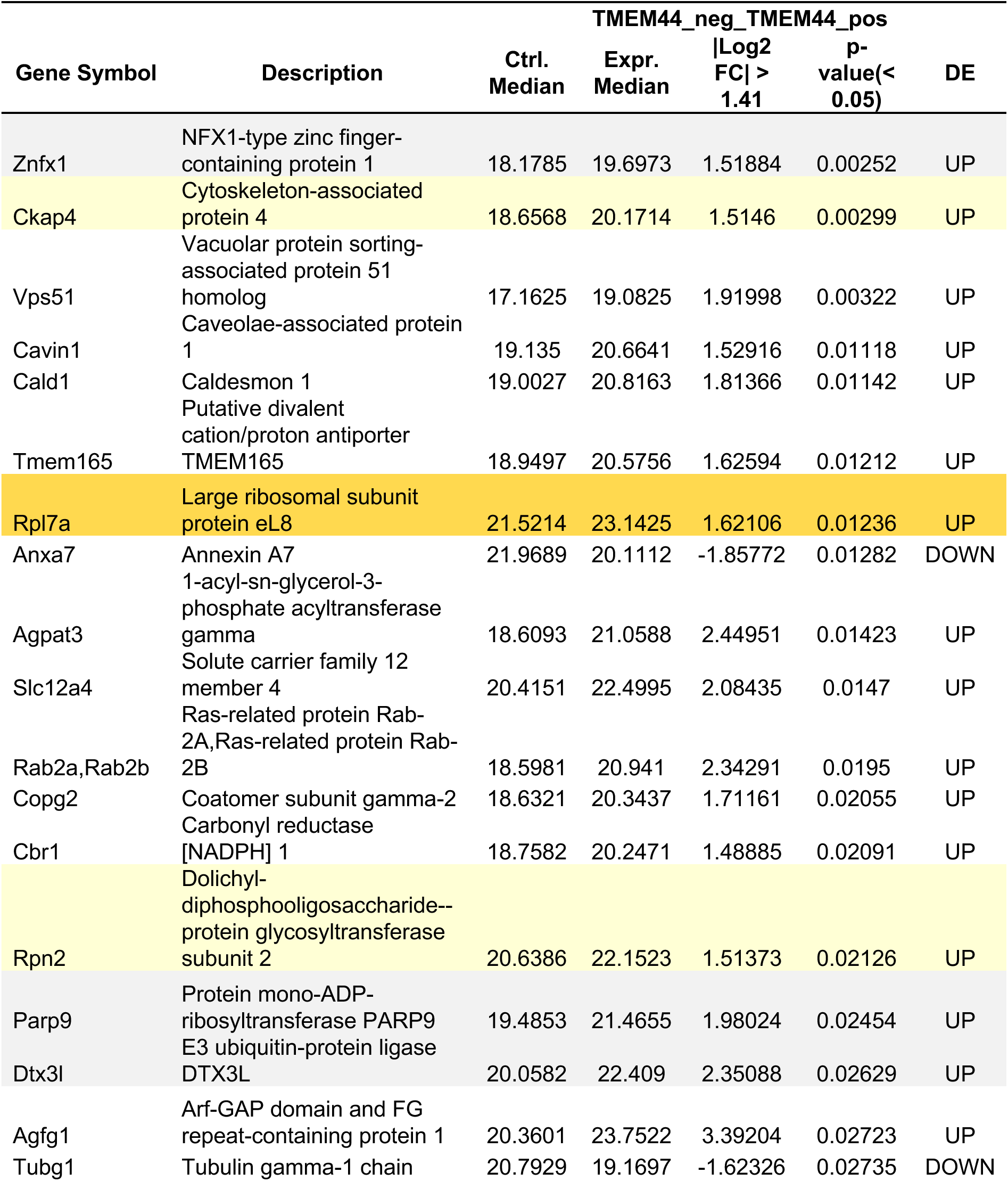
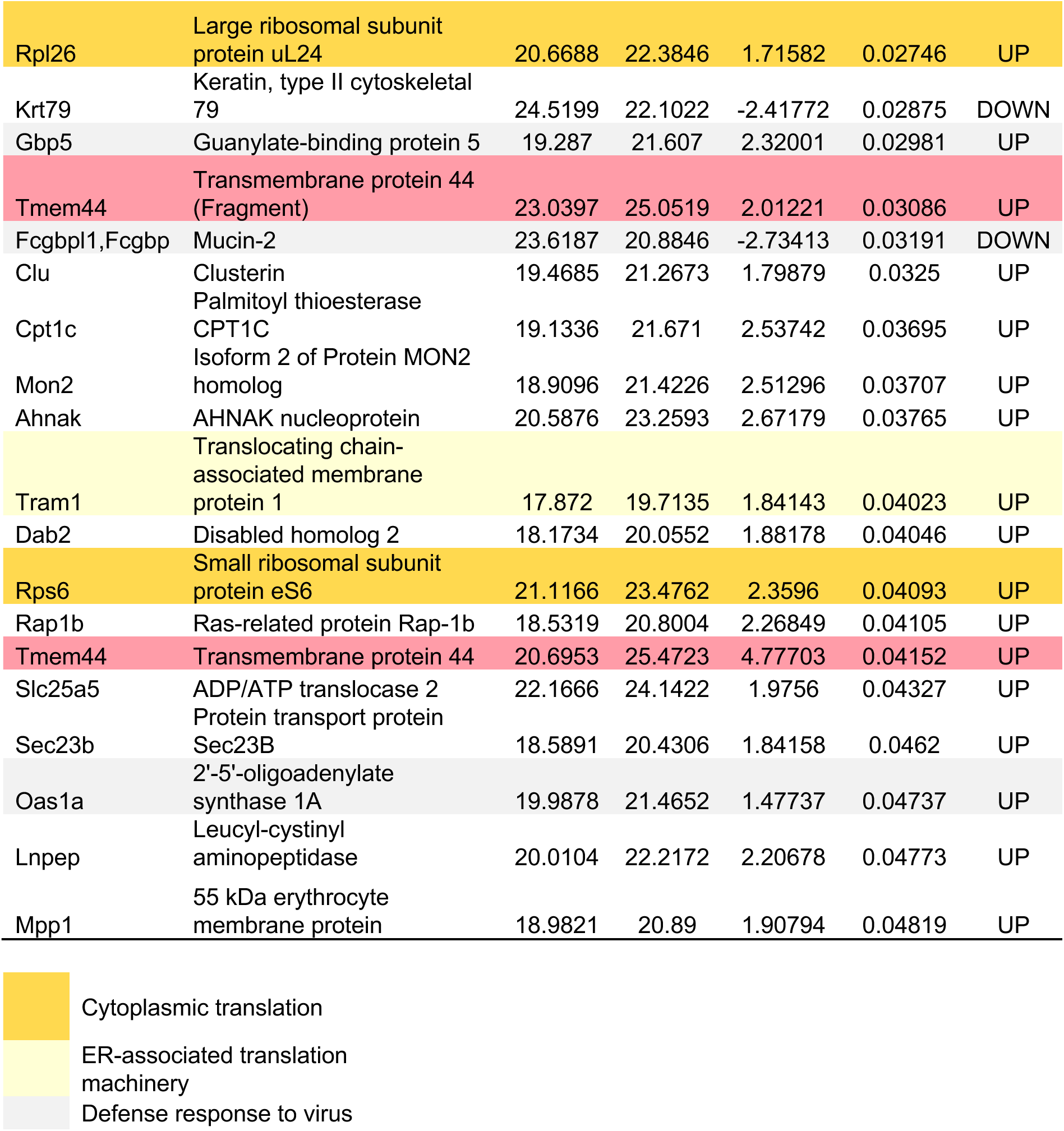
Differentially enriched proteins identified from TMEM44− versus TMEM44+ comparison. List of differentially enriched proteins identified from the TMEM44− versus TMEM44+ comparison. Yellow shading indicates proteins associated with cytoplasmic translation, light-yellow shading indicates endoplasmic reticulum–associated translation machinery components, and light-gray shading indicates proteins associated with defense response to virus.

**Table S3.**
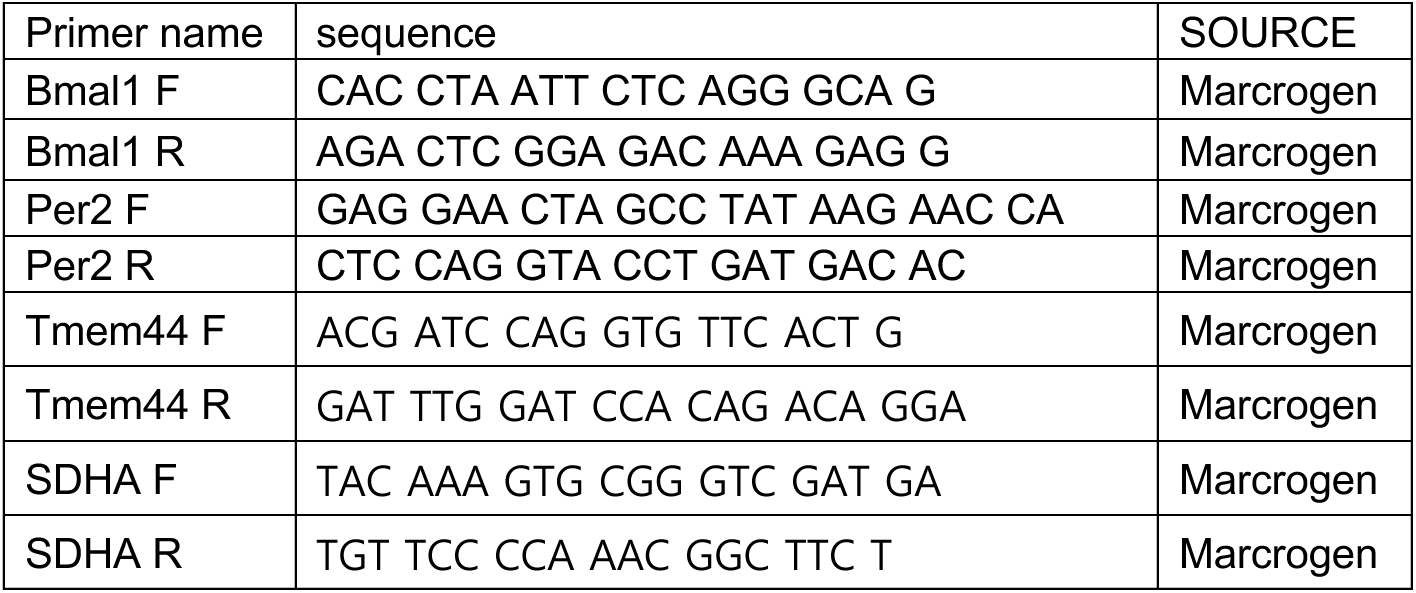
Primer sequences used for quantitative real-time PCR. List of primer sequences used for quantitative real-time PCR (qPCR) analysis in this study.

**Supplementary Figure 1.**
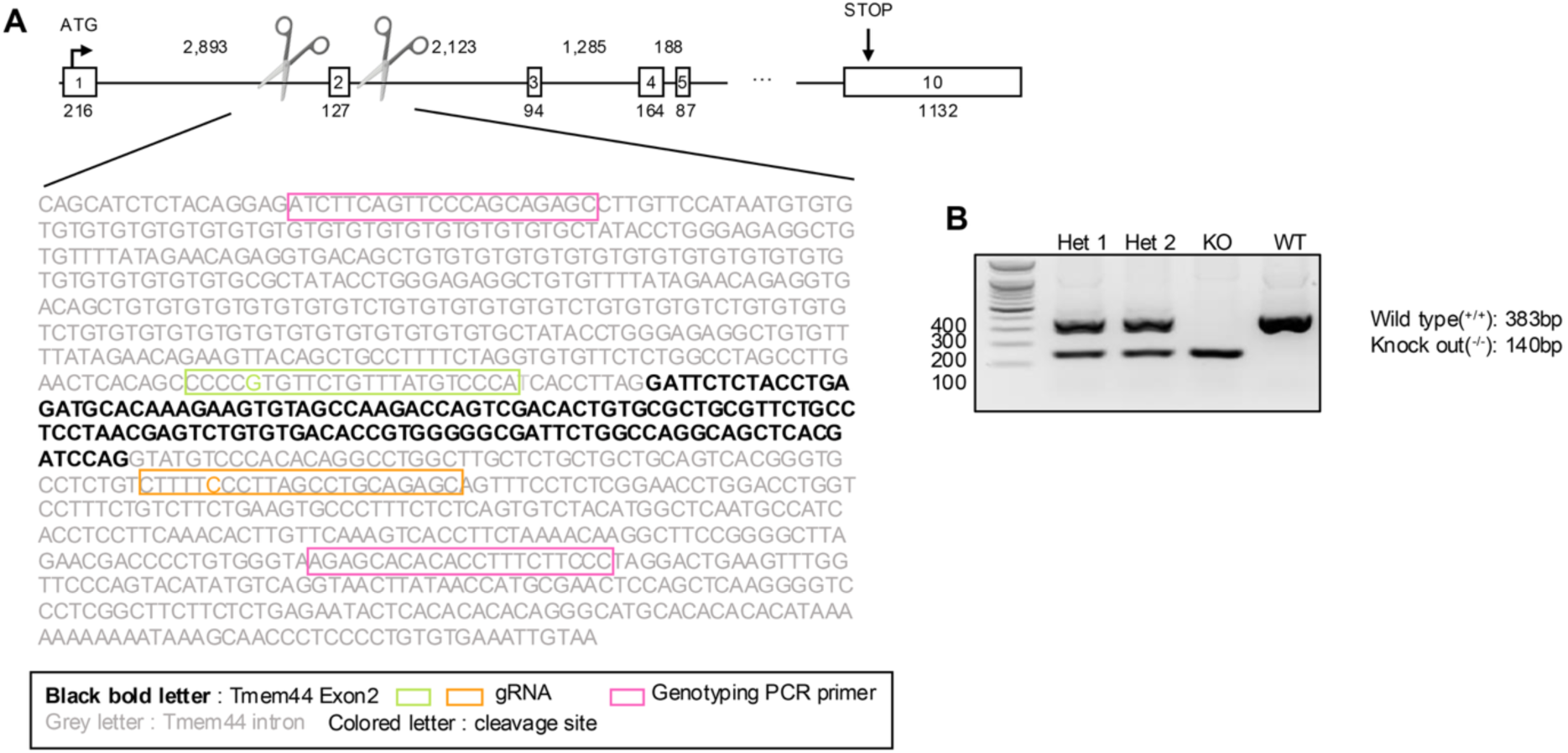
Generation of *Tmem44* knockout mice. (A) Schematic illustration of the *Tmem44* knockout strategy. Exon 2 was deleted using CRISPR–Cas9. The positions of gRNA target sequences, cleavage sites, and genotyping PCR primers are indicated. The corresponding genomic sequence is shown with exon regions and target sites highlighted. (B) Genotyping PCR analysis of *Tmem44*^+/+^, *Tmem44*^+/-^, and *Tmem44*^-/-^ mice.

**Supplementary Figure 2.**
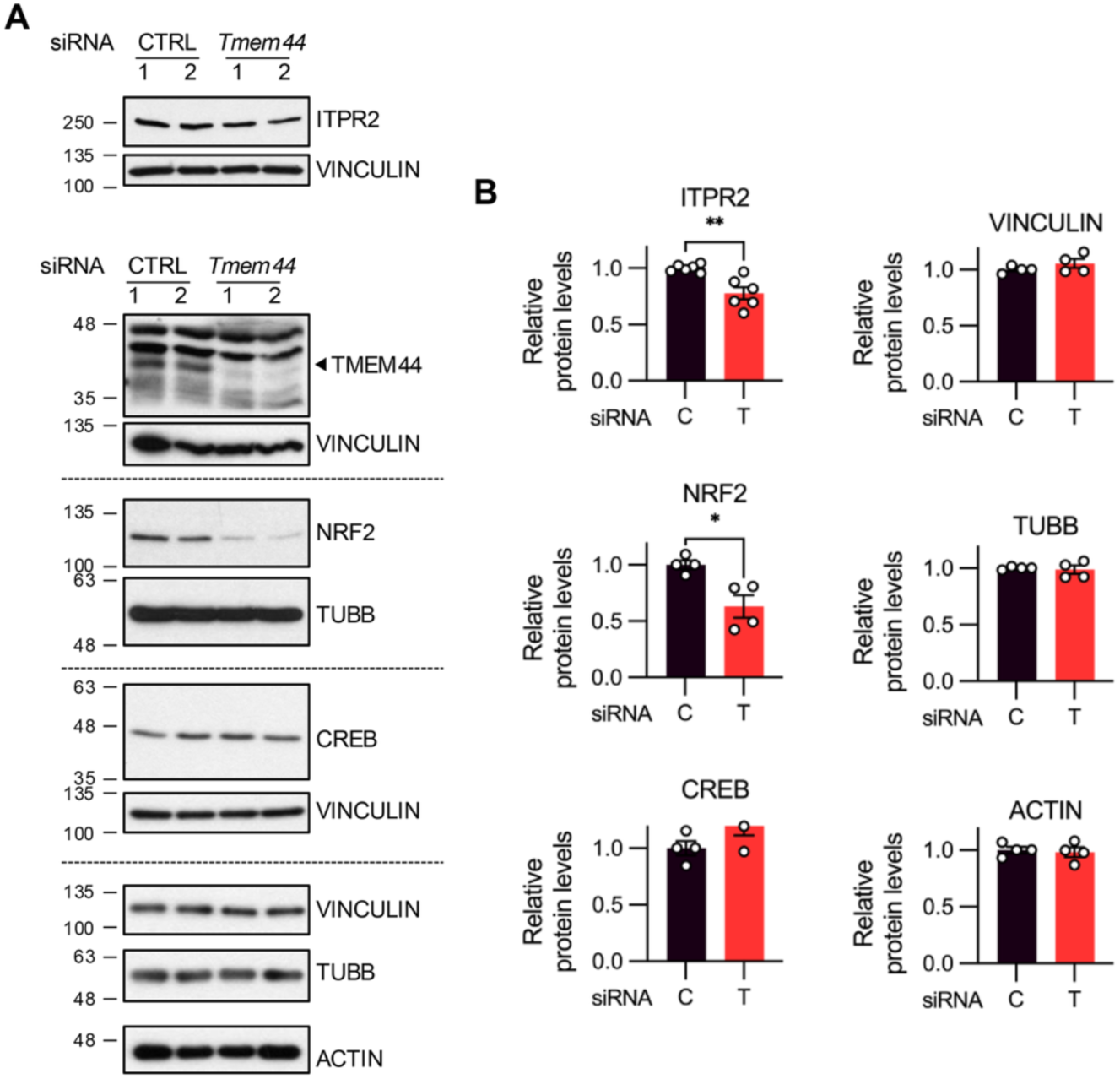
*Tmem44* knockdown does not uniformly reduce protein abundance in astrocytes. Astrocytes were analyzed 48 h after transfection with control or *Tmem44*-targeting siRNA. (A) Representative western blot images showing protein levels following *Tmem44* knockdown. (B) Densitometric analysis of protein levels from (A). NRF2 was normalized to TUBB, whereas ITPR2 and CREB were normalized to VINCULIN. Cytoskeletal proteins (VINCULIN, TUBB, and ACTIN) were not further normalized. All values were normalized to the mean of control astrocytes (set to 1). Control siRNA–treated astrocytes and *Tmem44* siRNA–treated astrocytes are labeled as C and T, respectively. Data are presented as mean ± SEM (ITPR2: n = 6; others: n = 4). Statistical significance was determined by Student’s t-test (*p < 0.05, **p < 0.01).

**Supplementary Figure 3.**
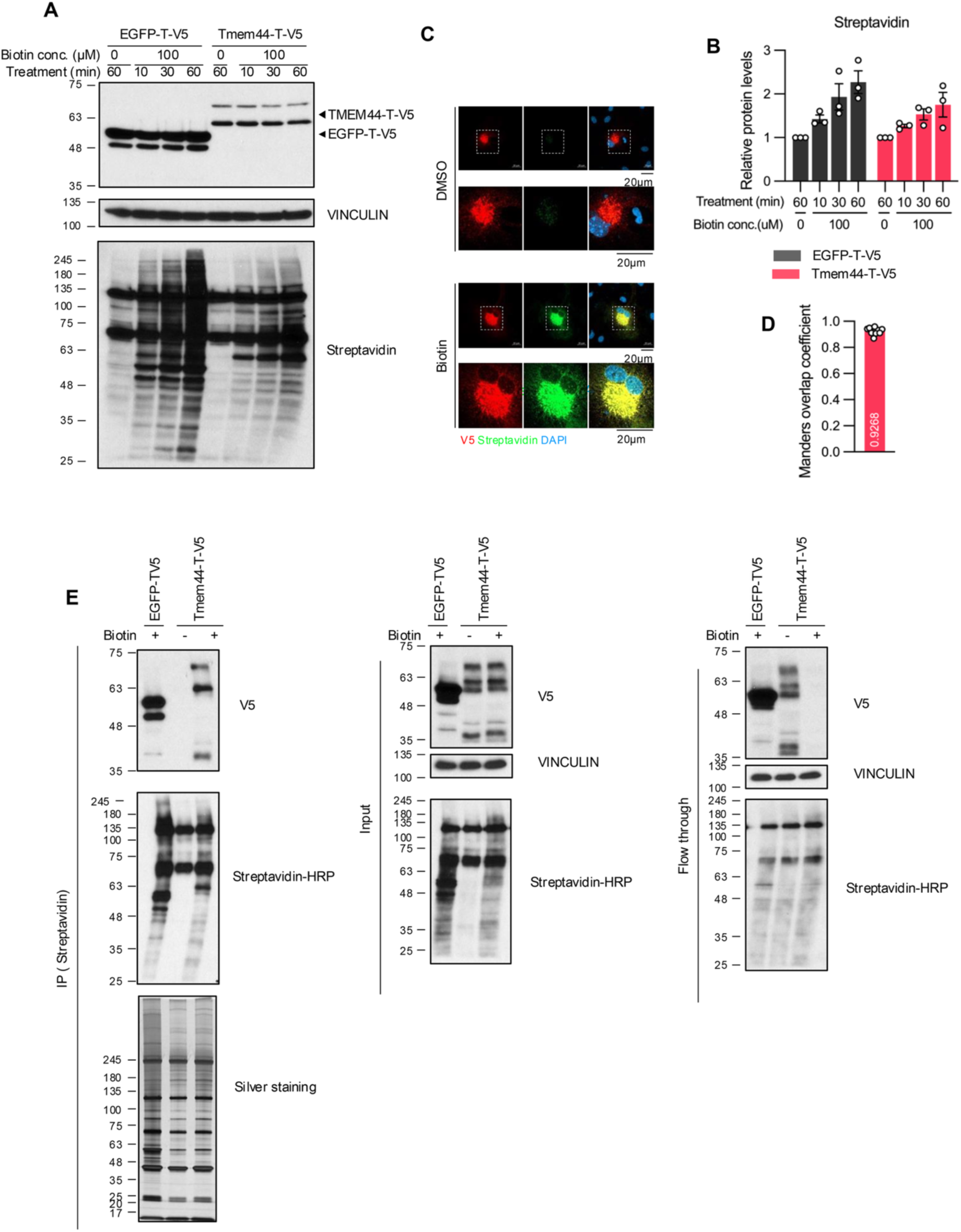
Optimization and validation of biotin labeling for miniTurbo-based proximity labeling in astrocytes. (A, B, E) TMEM44–miniTurbo–V5 and EGFP–miniTurbo–V5 are abbreviated as TMEM44–T–V5 and EGFP–T–V5, respectively, in the figure. Astrocytes were analyzed 48 h after transfection. (A) Representative western blot images showing biotinylated proteins under different biotin treatment durations. (B) Densitometric analysis of biotinylated protein levels from (A). Data are presented as mean ± SEM (n=3). (C) Representative immunocytochemistry images showing the subcellular localization of biotinylated proteins following treatment with DMSO or biotin (100 μM, 60 min). (D) Manders overlap coefficient analysis of colocalization shown in (n=11)(C). (E) Representative western blot images of streptavidin pull-down samples prior to proteomic analysis, showing input, pull-down, and flow-through fractions.

